# An Improved Systematic Method for Constructing Enzyme-Constrained Genome-Scale Metabolic Models Using a Protein-Chemical Transformer

**DOI:** 10.1101/2025.10.29.684458

**Authors:** Anna Schooneveld, Shafiat Dewan, Navot Arad, Sam Genway, Sonya Kaur Kalsi, Kotryna Bloznelyte, Will Addison, David Berman

## Abstract

Enzyme-constrained genome-scale metabolic models (ecGEMs) have improved Flux Balance Analysis (FBA) by incorporating enzyme turnover numbers (*k*_*cat*_s). Since in-vivo *k*_*cat*_ data is costly to obtain and therefore scarce, we present a novel multi-modal transformer-based approach with cross-attention to predict *k*_*cat*_ values for *Escherichia coli* using enzyme amino acid sequences and SMILES annotations of reaction substrates. For heteromeric enzymes, we evaluate multiple subunit *k*_*cat*_ aggregation strategies. We benchmark ecGEMs constructed with these strategies against current state-of-the-art models using experimental growth rates, ^13^C fluxes, and enzyme abundances, and prior to any calibration outperform or match existing methods. We also devise a new calibration method using flux control coefficients (derivatives of log flux with respect to log *k*_*cat*_), which we show to be identical to enzyme cost at the FBA optimum. Using these coefficients, we identify 8 key *k*_*cat*_ values to recalibrate using experimental data, subsequently achieving superior performance to the current state-of-the-art with 81% fewer calibrations.

## 1. Introduction

In recent decades, systems biology has progressed significantly with the development of genome-scale metabolic models (GEMs; Feist & Palsson 2008; Oberhardt et al. 2009; Gu et al. 2019). These models describe the interplay between genes, metabolites, and reactions in a mathematical framework that can be solved via Flux Balance Analysis (FBA; Orth et al. 2010). However, this approach is not sophisticated enough to capture all the behaviour of real cells. For example, *E. coli* exhibits a lower growth rate than predicted by simple FBA (Mao et al., 2022). One of the most promising solutions to this challenge has come from enzyme-constrained models, such as GECKO (Sánchez et al., 2017; Domenzain et al., 2022; Chen et al., 2024), sMOMENT/AutoPACMEN (Bekiaris & Klamt, 2020), and ECMpy (Mao et al., 2022; 2024). These models account for the fact that a cell has limited chemical resources for enzyme production, which constrains enzyme abundances and consequently the flux solution space, leading to more accurate predictions (Mao et al., 2022; Massaiu et al., 2019).

The quality of any enzyme-constrained model is strongly dependent on the accuracy of its enzyme turnover rates (Sánchez et al., 2017; Li et al., 2022). The turnover rate, *k*_*cat*_, of an enzyme is the number of substrate molecules catalysed by the enzyme per unit time. The literature can broadly be divided into two methods for obtaining *k*_*cat*_ values. Firstly, *k*_*cat*_s can be retrieved from experimental databases such as BRENDA (Chang et al., 2021) and SABIO-RK (Wittig et al., 2018). This is the approach taken by GECKO, AutoPACMEN, and ECMpy v2. The main issue with this method is that the *k*_*cat*_ coverage in these databases is incomplete even for well-studied organisms^1^, and very sparse or non-existent for less well-studied ones. Furthermore, measurements of *k*_*cat*_ values depend on experimental conditions, leading to non-uniformity between data from different studies.

With these issues in mind, a second approach uses machine learning to obtain *k*_*cat*_s. Examples of this approach include Heckmann et al. (2018), used in ECMpy 1.0, and DLKcat (Li et al., 2022). The former employed various machine learning approaches, such as linear regression and deep learning, using features like flux and catalytic site information obtained from protein structures. DLKcat is more general, as it does not require such features, making it suitable for less well-studied organisms. Instead, it requires inputs of SMILES strings for substrates (processed via a CNN), and amino acid sequences for enzymes (processed via a GNN). In the current work, we expand on this approach, presenting a novel transformer-based method (Vaswani et al., 2023) that takes SMILES strings and amino acid sequences as input. We demonstrate that our *k*_*cat*_ predictions outperform the state-of-the-art in terms of accuracy.

We apply our *k*_*cat*_ predictor to iML1515, the gold standard GEM for E. coli (Monk et al., 2017), to construct an enzyme-constrained GEM (ecGEM). Since existing *k*_*cat*_ predictors (including ours) only support monomeric reactions and the literature lacks consensus on how to combine subunit *k*_*cat*_ predictions for multimers, we evaluate the effect of multiple aggregation strategies on model performance through various benchmarks. These benchmarks also demonstrate that our model performs competitively with state-of-the-art approaches even before calibration.

Given the strong dependence of ecGEM performance on *k*_*cat*_ accuracy, calibration in the form of post-processing is often used to adjust *k*_*cat*_ using in vitro data. Generally enzyme cost is used to select reactions for calibration (Mao et al., 2022; 2024). We propose a more general alternative based on perturbative *k*_*cat*_ sensitivity analysis, using flux control coefficients (Kacser, 1973) to identify the most influential *k*_*cat*_ values. Although first disseminated in 1973, and later republished in 1995 (Kacser et al., 1995), flux control coefficients and metabolic control analysis have not been widely adopted for calibrating ecGEMs with FBA. We demonstrate that flux control coefficients are equivalent to enzyme costs in the case of an optimal FBA solution obeying enzyme constraints, thus providing the link between metabolic control analysis and enzyme costs determined through ecFBA solutions. Using our flux-control-coefficient-based method, we calibrate our ecGEM with significantly fewer ad-hoc *k*_*cat*_ corrections than other methods. The resulting ecGEM is on par with, or better than than existing approaches (Mao et al., 2022; 2024; Chen et al., 2024).

## 2. *k*_*cat*_ Prediction Transformer

This section details the data, architecture, and training process used to develop the *k*_*cat*_ model. The aim was to create a model that could predict the *k*_*cat*_ value given SMILES strings of the substrates and amino acids of the enzyme that catalyses the reaction.

### 2.1. Model Architecture

At a high level, our model architecture broadly follows the structure of DLKcat. It consists of three sub-models: two generate embeddings from substrates (as SMILES strings) and enzymes (as amino acid sequences), and a third predicts *k*_*cat*_ using these embeddings. Figure 1 shows a schematic representation. We introduce two major changes from DLK-cat’s approach: using pre-trained transformer-based models for embeddings (DLKcat trained a GNN for substrates and CNN for proteins), and introducing a custom 2-way cross-attention mechanism in the third sub-model.

**Figure 1.**
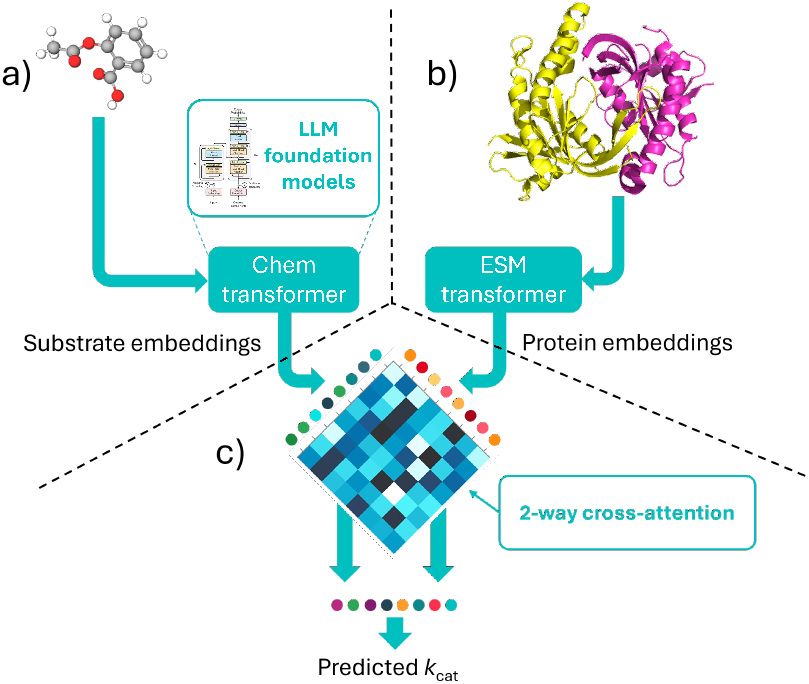
*k*_*cat*_ model architecture a) Pre-trained foundation model to generate molecule features. b) Pre-trained foundation model to generate protein features. c) 2-way cross-attention model to predict *k*_*cat*_ values

Our model requires two inputs. The first is a set of reaction substrates, each represented as a SMILES string (Weininger, 1988), concatenated into a single string using the non-bond token “.”, and passed to the Chem transformer (Chanda (2021); Figure 1a) to generate substrate embeddings. The second input is the enzyme’s amino acid sequence, which is passed to the ESM transformer (Lin et al., 2023) to produce protein embeddings. We removed the final projection layer from the Chem and ESM transformers, instead using their last hidden layers as input to the third sub-model, as this model requires embeddings (rather than the SMILES strings or amino acids output by the projection layer). For an input sequence of length *N* the generated embedding will be a matrix of size (*N, E*), where *E* is the fixed sized of the embedding.

The third sub-model implements a cross-attention mechanism between the substrate and protein embeddings produced by the Chem and ESM transformers, respectively. A complication with cross-attention is that it lacks order invariance, because the query (*Q*) matrix is derived from one input while the key (*K*) and value (*V* ) matrices are derived from the other, meaning the directionality of the attention (e.g., protein-to-substrate vs. substrate-to-protein) affects the output. Given that there is no natural “order” be-tween the a protein and its substrates, the model implements 2-way cross-attention (see Figure 2) which is invariant to permutations of the substrate and protein embeddings.

**Figure 2.**
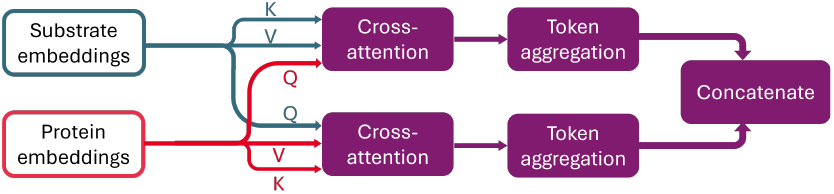
2-way cross-attention model architecture.

Following the cross-attention block in each branch, a token aggregation step is required to ensure that predictions are made over the complete sequence of the inputs, rather than individual tokens. We implemented three aggregation methods: (1) compute mean aggregation over the token embedding (2) take the embedding vector of the first token in the sequence (3) use a fuzzy membership function of a Gaussian mixture distribution and average memberships across the token dimension. Methods (1) and (2) produce an output vector of length *E*, (3) produces an output matrix with shape (*D, E*) where *D* is the number of 1-D distributions in the mixture. The choice of method is regarded as a hyperparameter.

The aggregated outputs from each branch are then concatenated and passed to the final projection layer to make the *k*_*cat*_ prediction (see Figure 2). All models were built using Python 3.9, PyTorch 1.13.1 (Paszke et al., 2019), and Lightning 2.0.3 (Falcon & The PyTorch Lightning team, 2019).

### 2.2 Data

To train the *k*_*cat*_ model, each data point must include a substrate set represented as SMILES strings, an enzyme given as an amino acid sequence, and a corresponding measured *k*_*cat*_ value. To obtain training data, measured *k*_*cat*_ values were collected from BRENDA (Chang et al., 2021) and SABIO-RK (Wittig et al., 2018). For reactions in BRENDA, substrate SMILES strings were retrieved programmatically from PubChem (Kim et al., 2024) via chemical name, and amino acid sequences from Uniprot by accession ID. For SABIO-RK, substrate SMILES strings were already present, and amino acid sequences were programmatically retrieved identically to BRENDA.

This training data has an issue of degeneracy at multiple levels. The first issue lies in molecular representation: a single molecule can have multiple valid SMILES strings. To address this, we standardised all SMILES strings by sanitizing with RDKit’s (Landrum et al., 2022) SanitizeMol function, removing isotopes, neutralizing charges, stripping stereochemistry, and converting to and from InChI to ensure tautomerism consistency. The second issue is that a single reaction can have multiple *k*_*cat*_ measurements. To resolve this, only the maximum *k*_*cat*_ is kept per unique reaction. Two data points are considered the same reaction if, after SMILES standardisation, they have identical substrates, products, and enzyme. The final degeneracy involves substrate ordering. Since reactions often have multiple substrates, their SMILES strings are concatenated before being input to the model. This introduces ambiguity in how to choose the order of concatenation. To address this, the dataset was augmented by including all possible permutations of substrate SMILES as distinct data points.

Before the final augmentation step, the training data was filtered to exclude entries with combined substrate SMILES over 510 tokens or amino acid sequences over 1000 characters, to accommodate the context limit of the Chem and ESM transformer respectively. To rebalance the data, *k*_*cat*_ values above 5000 were also discarded. This exluded only 1% of entries in the unfiltered dataset, which, despite spanning *k*_*cat*_ values of 0 − 10^8^, is heavily skewed towards smaller values.

The final dataset contained 35,499 entries, split into 65% train, 15% validation, and 20% test. Splitting was done prior to substrate permutation to avoid information leakage. It is important to note that, both the data and model architecture only support reactions catalysed by monomers; predictions for heteromers or homomultimers are not supported. The issue of how to deal with multimers will be revisited in Sections 4 and 3

### 2.3. Training and evaluation

During training, *k*_*cat*_ values are transformed via *y* = ln(*k*_*cat*_ + 1) to account for the logarithmic distribution of *k*_*cat*_s, and to prevent large values from dominating the loss. The addition of 1 ensures stability for small or zero values. As a result, the model also predicts in log-space (*ŷ*; see Figure 3), and the loss is computed between *y* and *ŷ*. The predicted value 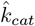 is retrieved by applying the inverse transformation 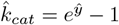.

**Figure 3.**
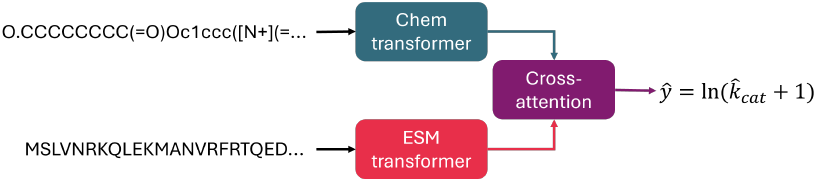
*k*_*cat*_ model inference data flow

Hyperparameter tuning was done using a Kubernetes cluster, with each hyperparameter set (a single model) trained on a single Nvidia V100 GPU. Every model was trained on the same training partition and evaluated on the validation set at the end of each epoch. The models were trained using the esm2 t33 650M UR50D model variant for the ESM transformer, Mean Squared Error (MSE) loss, and the Adam optimizer (Kingma & Ba, 2017).

The best model, as selected by the minimal validation loss, used mean token aggregation, a dropout of 0.2, 8 attention heads in each of the input branches of the cross-attention sub-model, a fixed learning rate of 10^−3^, and trained for 26 epochs with early stopping. This model took 24 hours to train. The selected model was evaluated on the held-out test partition, with 90% of predicted values being correct within 1 order of magnitude.

## 3. The Enzyme-Constrained Genome-Scale Metabolic Model

This section details the process of how our enzyme-constrained models were constructed. We constructed models for use with Python 3.10, COBRApy (Ebrahim et al., 2013).

### 3.1. Creating an enzyme-constrained model

As a base model, we use iML1515 (Monk et al., 2017), more specifically, the modified version in ECMpy (Mao et al., 2022), which corrects some minor known errors in iML1515. To make this model enzyme constrained, we follow and extend the approach in Mao et al. (2022).

To ensure that each enzyme-reaction pair has a unique *k*_*cat*_, we split reversible reactions into separate forward and backward reactions and split isoenzyme-catalysed reactions into different reactions. Unlike Mao et al. (2022), we add reverse reactions to all exchange reactions with a default lower bound of zero, to allow the intake of metabolites without requiring negative fluxes.

A GEM becomes an enzyme constraint model if, in addition to the standard FBA constraints,

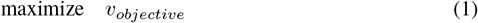

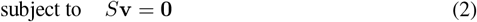

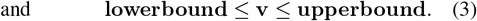

The model also obeys a total enzyme constraint

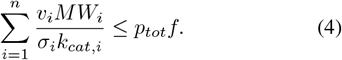

Here, *v*_*objective*_ is the objective flux (usually growth rate), **v** is a flux vector with elements *v*_*i*_ representing reactions (each with a lower and upper bound). *S* is the stoichiometric matrix, *n* is the total number of reactions, and *MW*_*i*_, *k*_*cat,i*_, and *σ*_*i*_ are the molecular weight, turnover number, and saturation coefficient of the enzyme catalysing a reaction. *p*_*tot*_ and *f* denote the total protein fraction and the enzyme mass fraction. Throughout this work, we assume *σ*_*r,i*_ = 1, consistent with (Mao et al., 2022; Bekiaris & Klamt, 2020). We set *p*_*tot*_ = 0.56 g gDW^−1^ based on experimental data (Bremer & Dennis, 2008; Brunk et al., 2016), and compute *f* as described in (Mao et al., 2022).^2^

A downside of the ECMpy approach is its incompatibility with major metabolic engineering packages like OptKnock (Burgard et al., 2003). Therefore, we adopt the method from Bekiaris & Klamt (2020), implementing the enzyme constraint via a pseudo-metabolite and pseudo-reaction. An “enzyme pool” pseudo-metabolite is added to each reaction *i* with a stoichiometric coefficient of 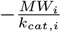. A pseudo-reaction is then added with this pseudo-metabolite as its only metabolite with an upper bound of *P* = *p*_*tot*_*f*. This is mathematically equivalent to Equation 4 (Bekiaris & Klamt, 2020). We then solved this system of equations via linear programming using COBRApy.

### 3.2. Annotations

To obtain the information we needed to implement the enzyme constraint (i.e. *MW*_*i*_ and *k*_*cat,i*_), we added additional reaction and metabolite annotations to our model. Amino acid sequences for reactions were obtained from the BiGG database, and SMILES strings for metabolites from MetaNetX. Deprecated MetaNetX annotations were manually updated using PubChem for some metabolites. Molecular weights and subunit information for genes were gathered from UniProt. As transport reactions require special treatment when setting *k*_*cat*_ values (see Section 3.3), we also annotated all transport reactions in the model. Appendix A provides more detail about the annotation process.

### 3.3. Determining *k*_*cat*_ values

Using these annotations, we passed amino acid sequences and SMILES strings for each non-transport reaction in our ecGEM to the *k*_*cat*_ predictor from Section 2. We did this for each enzyme subunit in the relevant reactions individually, predicting a *k*_*cat*_ value for each subunit. As monomers have only one subunit, this will be the final *k*_*cat*_ value. For multimeric reactions, i.e. those with multiple subunits, we came up with a strategy detailed below.

Our *k*_*cat*_ predictor does not produce meaningful predictions for transport reactions, as these are heavily under-represented in the training data. The BRENDA database contains *k*_*cat*_ values for only 31 out of 97 transport reactions in EC class 7 “Translocases”. The literature does not describe a principled method for handling transport *k*_*cat*_s. Therefore, we follow the common convention to use *k*_*cat*_ = 234, 000 *h*^−1^ (Corrao et al., 2024; Heckmann et al., 2018). Appendix B provides further details on the exact heuristics we used.

Reaction statistics and a brief discussion of these are provided in Table 3 and Appendix C. Our *k*_*cat*_ coverage is extensive - out of 5338 reactions with genes (the required input for our ML model), 5323 have a *k*_*cat*_ value.

#### 3.3.1. Calculating *k*_*cat*_ for Multimeric Enzymes

The literature on how to handle the choice of combining enzyme turnover rates for homomultimers and heteromers is sparse, and there is no clear consensus. ECMpy uses the *k*_*cat*_ with 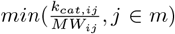, where *m* is the number of proteins in the complex (Mao et al., 2022), and DLKcat uses the maximum predicted *k*_*cat*_ value (Li et al., 2022). No rigorous derivation is given to explain why these heuristics were chosen.

However, rigorous handling of multimeric enzymes is crucial, as a non-neglicible portion of reactions are multimeric. In eciML1515, 46.5% of the 6191 reactions are homomultimeric, 32.5% are monomeric, 7% are heteromeric, and 14% are non-enzymatic (see Table 3). We address the calculation of *k*_*cat*_ for homomultimers and heteromultimers by evaluating models using various methods. For homomultimers comprised of *n* subunits, we calculate *k*_*cat*_ for the multimer as *n × k*_*cat*_ for the monomer^3^. For heteromers, we combine the *k*_*cat*_ value for each of the monomer/homomultimer present in the complex:

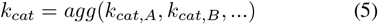

where *agg* is the minimum, maximum, average, or average weighted by the number of subunits. We produce one ecGEM for each aggregation method.

#### 3.3.2. *k*_*cat*_ Calibration targets via Flux Control Coefficients

It is inevitable that some *k*_*cat*_s are inaccurate due to machine learning errors or issues in handling multimers and transport reactions. To improve our enzyme-constrained model, we aim to select reactions for experimental calibration.

To understand which *k*_*cat*_ values most influence our ecGEM’s predictions, we quantified the sensitivity of target flux to individual enzyme turnover rates. Sensitivity is the magnitude of the derivative 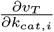, where *v*_*T*_ is the target flux and *i* is the enzyme-reaction pair. For each pair, we use FBA to solve for a target flux (e.g., biomass), apply a small perturbation 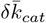 to 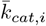( where 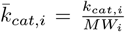), and re-solve the FBA model to compute the derivative as the ratio of flux change to 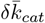. Finally, we rescale the derivative to obtain a dimensionless quantity:

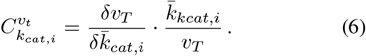

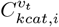 indicates the relative change in target flux when the *k*_*cat*_ of the *i*th reaction changes, known in metabolic control analysis as the flux control coefficient (Kacser, 1973; Kacser et al., 1995). A high flux control coefficient means the target flux is highly sensitive to that reaction. This analysis therefore identifies reactions that bottleneck the target flux. High 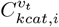 genes highlight the most efficient improvements to a metabolic model’s predictive power from lab measurements.

When measuring *k*_*cat*_ in vivo, this informs where to invest lab effort to manually calibrate and improve the model. This is useful for metabolic engineering, guiding the choice of proteins/enzymes to improve through protein engineering or identifying genes for upregulation via promoter engineering. Additionally, it benefits general model accuracy. The top *N k*_*cat*_ values from this analysis indicate the enzymes that best explain variance in model performance. This reduces the problem space from over 6,000 reactions to tens or hundreds. Combined with tools like metabolic flux visualizations (Figure 6, and 7), this helps identify key pathways that enhance model accuracy or target flux.

To our knowledge, perturbative *k*_*cat*_ analysis using flux control coefficients has not been previously applied to ecGEM calibration in the literature. ECMpy v2 (Mao et al., 2024) provide a related approach, but they iteratively calibrate based on enzyme cost 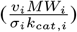. Notably, for an optimised ecGEM, the flux control coefficient is mathematically equivalent to enzyme cost. A detailed proof and further discussion are provided in Appendix E. A key advantage of using flux control coefficients is their generality: they will inform us of flux bottlenecks even when additional constraints are introduced to the model.

## 4. Results

### 4.1. Computing benchmarks

We captured the performance of our ecGEMs using a selection of benchmarks outlined below, and compared our models with reference models from the literature. Optimisation was performed via pFBA (Lewis et al., 2010) with CPLEX.

- **Glucose growth rate**. This benchmark compares the model growth rate on a glucose substrate to the experimental value (0.66 h^−1^ (Adadi et al., 2012)). It is given as a signed percentage difference.
- **Flux comparison to measured** ^**13**^**C fluxes (RMSE)**. This benchmark compares model fluxes to measured ^13^C fluxes (Okahashi et al., 2014) via the Root Mean Squared Error (RMSE). We sum isoenzyme-catalysed reactions and subtract reverse fluxes.
- **Growth rate on different substrates**. This benchmark compares the simulated growth rate on 24 different carbon sources to existing experimental measurements (Adadi et al., 2012), again using the RMSE.
- **Enzyme abundances**. This benchmark compares predicted enzyme abundances in our model to enzyme abundances from Corrao et al. (2024), via the root mean squared logarithmic error (RMSLE). Further detail about this abundance analysis is provided in Appendix F.

### 4.2. Models to evaluate

We produced four uncalibrated versions of eciML1515 (min, max, avg, wavg), one for each *k*_*cat*_ aggregation method from Section 3.3.1. We also create min 4 clbr and min 8 clbr, which are the min model where four or eight reactions are calibrated respectively (see Section 4.5 for more detail).

We benchmark against several state-of-the-art enzyme-constrained models from the literature (Mao et al., 2022; 2024). More detail on the specifics of these models can be found in Appendix G

- **ECMpy ML** is the ECMpy v1 model without any *k*_*cat*_ calibration; it uses machine learning-derived *k*_*cat*_ values from (Heckmann et al., 2018).
- **ECMpy expmnt** is the ECMpy v2 model without calibration, made up of experimental *k*_*cat*_ values from BRENDA and SABIO-RK. An average *k*_*cat*_ is used for reactions not in the databases.
- **ECMpy DLKcat** was created via the DLKcat pipeline in ECMpy v2. This pipeline produced NaNs for 15% of *k*_*cat*_s due to SMILES retrieval issues; we left those reactions unconstrained. Note therefore that results for this model reflect the combined ECMpy+DLKcat performance rather than DLKcat alone.
- **ECMpy ML clbr** is ECMpy ML after calibration, where 14 reactions with high enzyme cost were updated using experimental *k*_*cat*_s from BRENDA and SABIO-RK.
- **ECMpy expmnt clbr** is ECMpy expmnt after the v2 calibration process, which consists 50 rounds of iteratively replacing high enzyme cost reactions with the largest available experimental value from BRENDA or SABIO-RK.

### 4.3. Our *k*_*cat*_ predictor creates more accurate predictions of metabolism

We first run the benchmarking on all the uncalibrated models. The results are shown in Table 1. Without any calibration, the performance of our models is generally superior to the benchmark models. Our models are the highest scoring in terms of glucose growth, ^13^C, and substrate RMSE. The only area where a model from the literature is superior is for the abundance analysis, where ECMpy ML takes the top spot. The fact that our models are overall the best performing on most benchmarks suggests that our *k*_*cat*_ estimates are superior to the current state-of-the-art, and would provide a better basis for an ecGEM prior to any calibration.

**Table 1.** Benchmarking results of uncalibrated ecGEMs against experimental data from the literature for growth rates, ^13^C fluxes, and enzyme abundances. A description of how these metrics were computed can be found in Section 4.1. Glucose growth is given as a signed percentage difference between simulated and experimental growth rate. These models have not undergone any *k*_*cat*_ calibration, see Section 4.2 and Appendix G for details on how these models were obtained. The best performing result for each benchmark is indicated in **bold**.

| ecGEM | Glucose growth % error | $^{13}\text{C}$ RMSE | Various substrate growth RMSE | Abundance RMSLE glucose $\times 10^{-3}$ |
| --- | --- | --- | --- | --- |
| min | -57.2 | 6.18 | 0.31 | 6.67 |
| max | 29.4 | **2.47** | 0.31 | 6.50 |
| avg | **12.3** | 2.70 | 0.22 | 4.58 |
| wavg | -14.3 | 9.17 | **0.13** | 6.16 |
| ECMpy_ML | -44.2 | 4.12 | 0.20 | **3.41** |
| ECMpy_exp | -73.7 | 4.42 | 0.34 | 6.93 |
| ECMpy_DLKcat | -45.4 | 5.13 | 0.24 | 6.45 |

Note also that overall, our ecGEMs outperform the ECMpy DLKcat model. This confirms that the improved performance of our *k*_*cat*_ predictor over DLKcat as noted in Figure 4 also translates to measurable improvements in model performance at the ecGEM scale.

**Figure 4.**
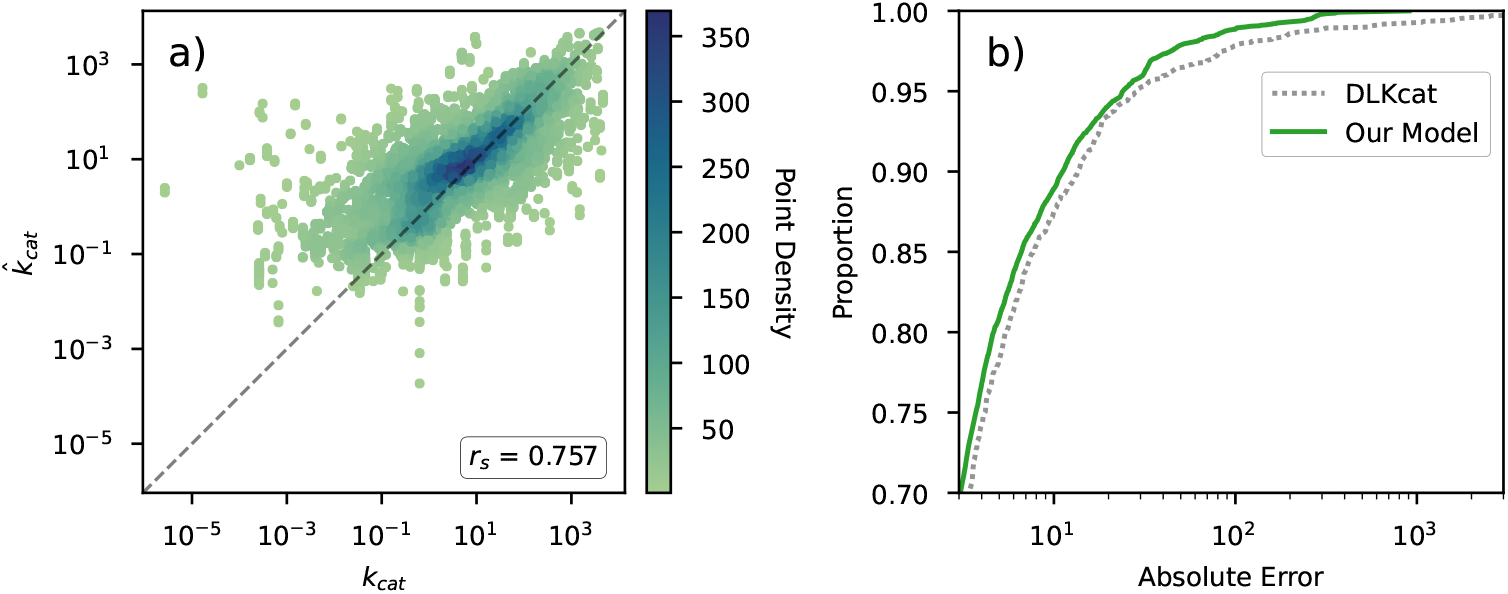
*k*_*cat*_ model performance. a) log-log scatter plot showing measured *k*_*cat*_ vs predicted 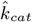. Spearman’s rank correlation coefficient *r*_*s*_ = 0.757 for our model, compared to *r*_*s*_ = 0.720 for DLKcat. b) Comparison of the error in model predictions between our model and DLKcat. At each point on the x-axis, the curves show the proportion of test set predictions with the same cumulative error or less. Our model makes notably fewer predictions that have large errors.

### 4.4. Uncovering sensitivity to modelling of heteromers

Our ecGEM performance also varies substantially depending on the method we use for aggregating *k*_*cat*_ predictions for heteromers, and other than the fact that the min model generally performs worst, there is no one method that is clearly superior to the others. This is an important finding, as it shows that the choice of aggregation strategy can have a considerable impact on ecGEM quality. As described in Section 3.3.1, the current literature lacks a cohesive and considered approach to this decision. However, our results imply that it is an important factor for ecGEM performance. We therefore strongly recommend more research into this issue in order for machine learning *k*_*cat*_s to reach their full potential.

### 4.5. Model improvement with fewer calibrations

As our min model had the worst growth rate error on glucose (57.2%), we selected this model for calibration. Using our method from Section 3.3.2, we computed the flux control coefficients 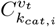 (see Figure 8) to identify the top 10 most sensitive reactions as calibration targets to improve (See Table 4). Ideally, we would have been able to measure *k*_*cat*_ for these 10 reactions in vivo, but we did not have the required laboratory resources. Instead, consistent with Mao et al. (2022; 2024), we replaced the *k*_*cat*_ value with the highest reported *k*_*cat*_ from BRENDA (Chang et al., 2021) and SABIO-RK (Wittig et al., 2018) via the EC number(s).

To quantify how many reactions we would need to adjust to achieve improvements in ecGEM performance, we created ecGEMs with between 1 and 10 *k*_*cat*_ values calibrated, and computed their glucose growth rate error values. Figure 5 shows the improvement in glucose growth rate error when successive *k*_*cat*_ values (ranked by *k*_*cat*_ sensitivity) are adjusted cumulatively. Substantial improvements are made with just 4 reactions adjusted, and the improvement plateaus after just 8. We compare these two models (min 4 clbr and min 8 clbr) to similar models from the literature in Table 2.

**Table 2.**
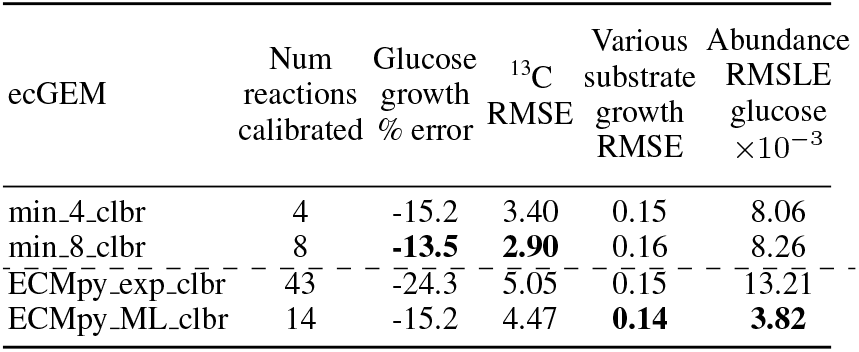
Benchmarking results of calibrated ecGEMs against experimental data from the literature for growth rates, ^13^C fluxes, and enzyme abundances. A description of how these metrics were computed can be found in Section 4.1. Glucose growth is given as a percentage difference between simulated and experimental growth rate. These models have been optimised with *k*_*cat*_ calibration, see Section 4.2 for details on how the ECMpy models were obtained, and Section 4.5 for the creation of the min 4 clbr and min 8 clbr models. The best performing result for each benchmark is indicated in **bold**.

**Figure 5.**
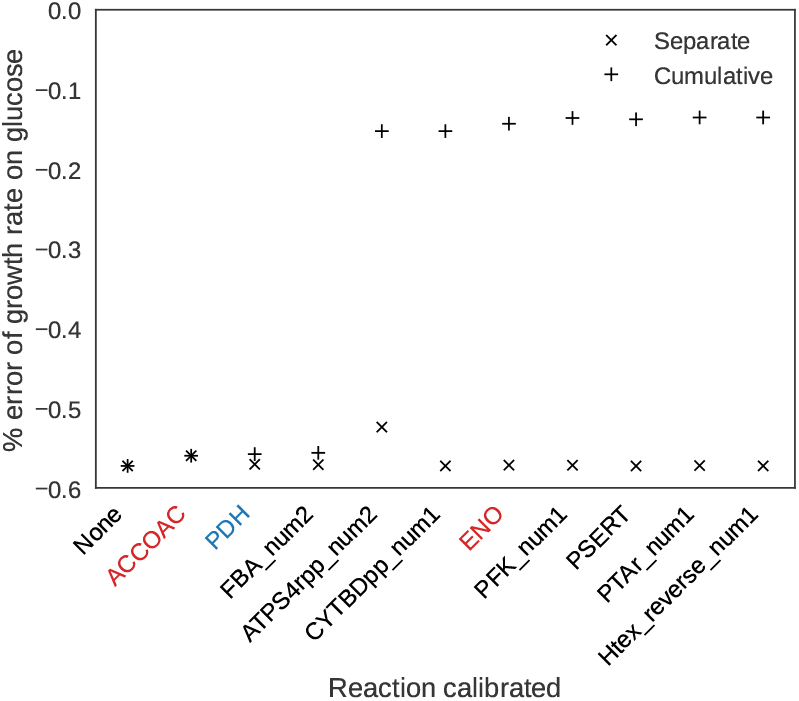
Calibrated min model with reactions calibrated separately and cumulatively (left to right). Red highlighting corresponds to ECMpy v1 round 1 enzyme abundance corrections, Blue highlighting corresponds to ECMpy v1 round 2 ^13^C corrections.

Calibration substantially improves our min model across most metrics — adjusting just four reactions yields comparable performance to ECMpy ML clbr, which calibrated 14 *k*_*cat*_s. Increasing the number of calibrated reactions to eight leads to the lowest growth rate error among all calibrated models, in fewer calibrations than the benchmark models. This model achieves an reduction in glucose growth-rate error from -57.2% to -13.5% (Figure 5) and RMSE on ^13^C flux data from 6.18 to 2.90 (Tables 1, 2). The significant reduction in growth rate error only occurs when calibrated *k*_*cat*_ values are applied cumulatively^4^, not separately, as shown in see Figure 5.

We identify ATPS4rpp num2 as a major biomass flux bottleneck - calibrating its *k*_*cat*_ reduces the glucose growth rate error by 40%. Our predicted *k*_*cat*_ differs from the experimental value in BRENDA by two orders of magnitude (Table 4). The reaction is catalysed by an eight-subunit enzyme complex (Moore et al., 2024). Our *k*_*cat*_ predictor, trained only on monomers, outputs values spanning four orders of magnitude, and for heteromers, our simple aggregation method is highly sensitive to outliers. Notably, 18 of the top 100 reactions ranked by 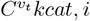 are heteromeric (Figure 9). This is a substantial amount, given that only 7% of all reactions in our ecGEM are heteromeric (Table 3). For all multimers combined (including homomultimers), 81 top 100 reactions are multimeric, compared to 54% of all ecGEM reactions, Together, all these observations highlight the need for a *k*_*cat*_ predictor that can directly estimate turnover numbers for heteromeric complexes, and multimers in general.

**Table 3.** Reaction statistics for eciML1515.

| Metric | Value | % of Total |
| --- | --- | --- |
| Total reactions | 6190 | - |
| Total reactions with genes | 5338 | 86% |
| Coverage of $k_{cat}$ incl transport | 5323 | 86% |
| Missing $k_{cat}$ | 15 | 0.2 % |
| ML $k_{cat}$ | 2360 | 38% |
| Exchange reactions (EX<sub>-</sub>) | 662 | 11% |
| Spontaneous reactions (s0001) | 64 | 1% |
| Transport reactions | 2963 | 48% |
| Porin reactions | 2186 | 35% |
| Monomers | 2020 | 33% |
| Homomultimers | 2883 | 47% |
| Heteromultimers | 435 | 7% |

Our calibration targets for the min model partially overlap with ECMpy v1 (Mao et al., 2022), sharing two round 1 enzyme usage corrections and one round 2 ^13^C flux correction. However, we need just 4 calibrations (Figure 5) to reduce glucose growth rate error to 15% whereas ECMpy v1 uses 14 and ECMpy v2 uses 43. Given that our method is mathematically equivalent to theirs (as shown in Appendix E), the fact that we need fewer calibrations implies that our initial *k*_*cat*_ estimates must be more accurate, highlighting the accuracy of our machine learning predictor.

## 5. Conclusion

Using the novel transformer-based *k*_*cat*_ predictor introduced in this work, we produced an enzyme constrained genome scale model which outperforms the current state-of-the-art for ecGEMs that have not been calibrated using lab data. Furthermore, with our *k*_*cat*_ sensitivity analysis, we have devised a rigorous way to identify which *k*_*cat*_s would benefit the most from calibration with lab data. When applied to the min model, our worst performing uncalibrated model, this process makes our model competitive with the best calibrated models in the literature, but by calibrating 81% fewer *k*_*cat*_ values. Thus, our findings add to the growing body of evidence (Li et al., 2022; Heckmann et al., 2018) that machine learning for predicting turnover numbers is a core part of future systems biology.

However, to produce optimal machine learning *k*_*cat*_s, we have found that it is important to have a well founded method for estimating *k*_*cat*_s for multimers. This is currently an under-explored area of study, and different research groups adopt different methods without a clear justification. Our findings show that ecGEM performance can vary significantly depending on the method used for aggregating *k*_*cat*_ values for heteromeric enzymes, and this choice therefore warrants further research. We emphasise that any machine learning predictor for enzyme turnover rates for heteromers and homomultimers needs to be more advanced than the currently available methods. It is likely that a single uniform aggregation method across all reactions in a given model is too simplistic. The function of subunits and the way in which they combine to form an enzyme complex varies significantly from enzyme to enzyme, and it seems biologically likely that the optimal aggregation method might differ per enzyme and therefore require a more complex method than a simple max, min, avg, or wavg operation. Multimeric reactions make up 81 of the top 100 most senstive reactions (Figure 9), so getting these *k*_*cat*_s right is crucial.

Therefore, to imbibe as much biological context as possible, an improved future *k*_*cat*_ predictor should be trained on a dataset that includes many more examples of reactions catalysed by homomultimers and heteromers and have an architecture that supports multimeric inputs. A second direction of improvement would center around enhanced capabilities for dealing with transport and exchange reactions. To do this, the model would have to encounter a substantial number of these reactions during training.

However, no matter how advanced the method for estimating *k*_*cat*_, that there will be inaccuracies in estimates/predictions, whether from deep learning or otherwise. So, in order to refine an ecGEM, there needs to be a rigorous calibration method used. We have shown that the method of calibration via per-reaction enzyme cost calculations is mathematically equivalent to calculating the flux control coefficient per reaction. Furthermore, we don’t expect this equivalence to hold when additional constraints are added to the optimisation. However our method of calculating the flux control coefficient through perturbing individual *k*_*cat*_ will generalise to any ecGEM.

## Acknowledgements

We thank the reviewers for their feedback on the submitted version of the manuscript. We are grateful to Jakub Chromý, Luca Rossoni, and Ian Taylor for their expert input on metabolic engineering. We also thank Sadhira Wagiswara for creating the visualisations of our metabolic networks. Finally, we thank Chris Roberts, Frances Metcalfe, Richard Snell, Sally Epstein and those at Capgemini Invent for sponsoring and supporting this project and the subsequent publishing effort.

## Impact Statement

Improved metabolic modelling of organisms like *Escherichia coli* requires information about enzyme dynamics. We present a novel, transformer-based machine learning architecture with two-way cross attention to predict enzyme turnover numbers to present a state-of-the-art method for constructing metabolic models. Our insights will provide a deeper understanding of constructing and evaluating metabolic models in the field of Metabolic Engineering.

## A. ecGEM Annotations

In order to obtain the information we need to implement the enzyme constraint (i.e. *MW*_*i*_ and *k*_*cat,i*_), we add additional reaction and metabolite annotations to our model file. We extracted amino acid sequence annotations from the BiGG (King et al., 2015) database to annotate reactions. We annotated metabolites with SMILES strings extracted from the MetaNetX database (Moretti et al., 2020; 2015; Ganter et al., 2013; Bernard et al., 2012). It must be noted that a lot of the pre-existing MetaNetX annotations are deprecated for both the ECMpy eciML1515 model and the BiGG database, such as prbatp c (MNXM1351). For the majority of metabolites, this wasn’t an issue, since the deprecated reference had syntactically correct SMILES annotations associated. However, for other reactions, such as octapb c (MNXM147531), we had to manually fill these in using the canonical SMILES from PubChem (Kim et al., 2024). We did this for 139 out of 1877 metabolites. Finally, we used UniProt (Consortium, 2022) to obtain molecular weights and subunit information associated with the genes in *E. coli*. Together, these three annotation sets were required to predict *k*_*cat*_s: the first two for the machine learning model input, and the last for the aggregation stage for multimer/heteromer *k*_*cat*_ prediction.

Given the necessity for special treatment of *k*_*cat*_ values for transport reactions (see Section 3.3), we also annotated all transport reactions in the model according to their classification in (Heckmann et al., 2020). We discovered that some transport reactions were not in this list, so we decided to add an annotation of “Transport, uncategorised” to all reactions that have either “transport” or “exchange” in the name and that do not have either a gene reaction rule of “s0001” (spontaneous reactions) or “” (empty gene^5^). The latter two types were not annotated as transport reactions in our model.

## B. Transport Reactions

Because our *k*_*cat*_ predictor cannot predict transport reactions well (due to the fact that they are vastly underrepresented in the training data), and there is no principled method for handling transport turnover numbers, we followed the common convention to use *k*_*cat*_ = 234, 000 *h*^−1^ (Corrao et al., 2024; Heckmann et al., 2018). This choice is somewhat arbitrary, and we recommend future research to find a more rigorous treatment. The precise heuristics we used are the following:

- For all transport reactions that are annotated as a transport reaction but are not porins, e.g. SUCptspp, we set *k*_*cat*_ = 234, 000 h^−1^.
- For transport reactions with one pore, e.g. ACtex gene b1377 (ompN), we also set *k*_*cat*_ = 234, 000 h^−1^.
- For transport reactions with N pores e.g. ACtex gene b0241 (phoE), we set *k*_*cat*_ = *N ×* 234, 000 h^−1^ (*N* = 3 for phoE).
- For spontaneous reactions with gene s0001 or reactions without genes, we do not set *k*_*cat*_, which means these reactions are unconstrained.

## C. ecGEM metrics

Table 3 shows the reaction statistics for our model. *k*_*cat*_ coverage in our ecGEM is extensive; we have *k*_*cat*_ values for 5323 out of 5338 reactions for which the required input information (genes) is available. For 7 of the 15 missing reactions, the model output a *k*_*cat*_ of 0. The remaining 8 reactions have too long substrate SMILES strings, exceeding the ML model’s 512 substrate token limit. These 15 reactions (only 0.2% of all reactions) were left unconstrained. The rest are constrained by either our ML *k*_*cat*_s or the transport values in Appendix B.

In terms of subunits, the largest group consists of homomultimers, which make up 47% of all reactions. 7% of reactions are heteromeric, for which the choice of multimer aggregation described in Section 3.3.1 affects the *k*_*cat*_ value, it is important to note that this is a non-negligible number.

## D. Flux visualisations

Figures 6 and 7 show flux visualisations for the min model. Edges correspond to ln *v*_*i*_. Where ln *v*_*i*_ *<* 0.5, and inclusion significantly reduces visual clarity, nodes and edges are not shown. Grey nodes and edges in Figure 6 and 7 correspond to flux solutions which fall below the threshold. Key:

**Figure 6.**
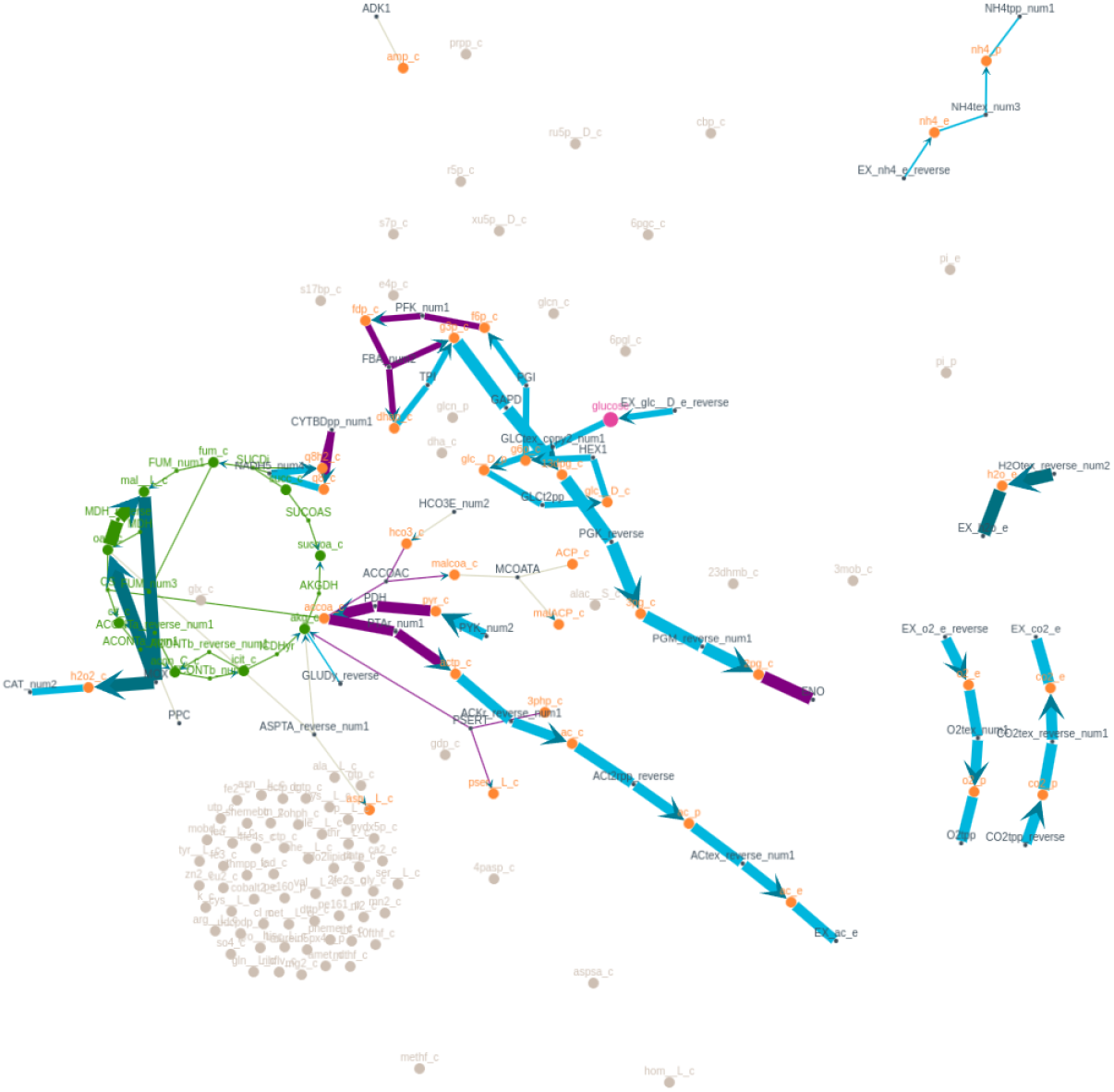
Metabolic Flux Visualisation: Enzyme-constrained solution, found with FBA for the min model.

|  |  |
| --- | --- |
| **Green** | Citric Acid Cycle |
| **Purple** | Calibration candidates (Section 4.5) |
| **Pink** | Glucose |
| **Grey Nodes** | Metabolites of reactions falling below flux threshold |
| **Grey Edges** | Fluxes clamped at minimum value for visual clarity (0.5) |
| **Dark Blue** | Fluxes clamped at maximum value for visual clarity (10.0) |
| **Orange** | Other Metabolites |
| **Light Blue** | Other Reactions |

Figure 6 shows the fluxes in the min model before any *k*_*cat*_ calibration was applied. Figure 7 shows the fluxes after calibrating the 8 most sensitive reactions via our flux control analysis. When visually comparing these two, it is clear that *k*_*cat*_ calibration of a relatively small number of reactions can have an effect on the whole metabolism of the organism.

**Figure 7.**
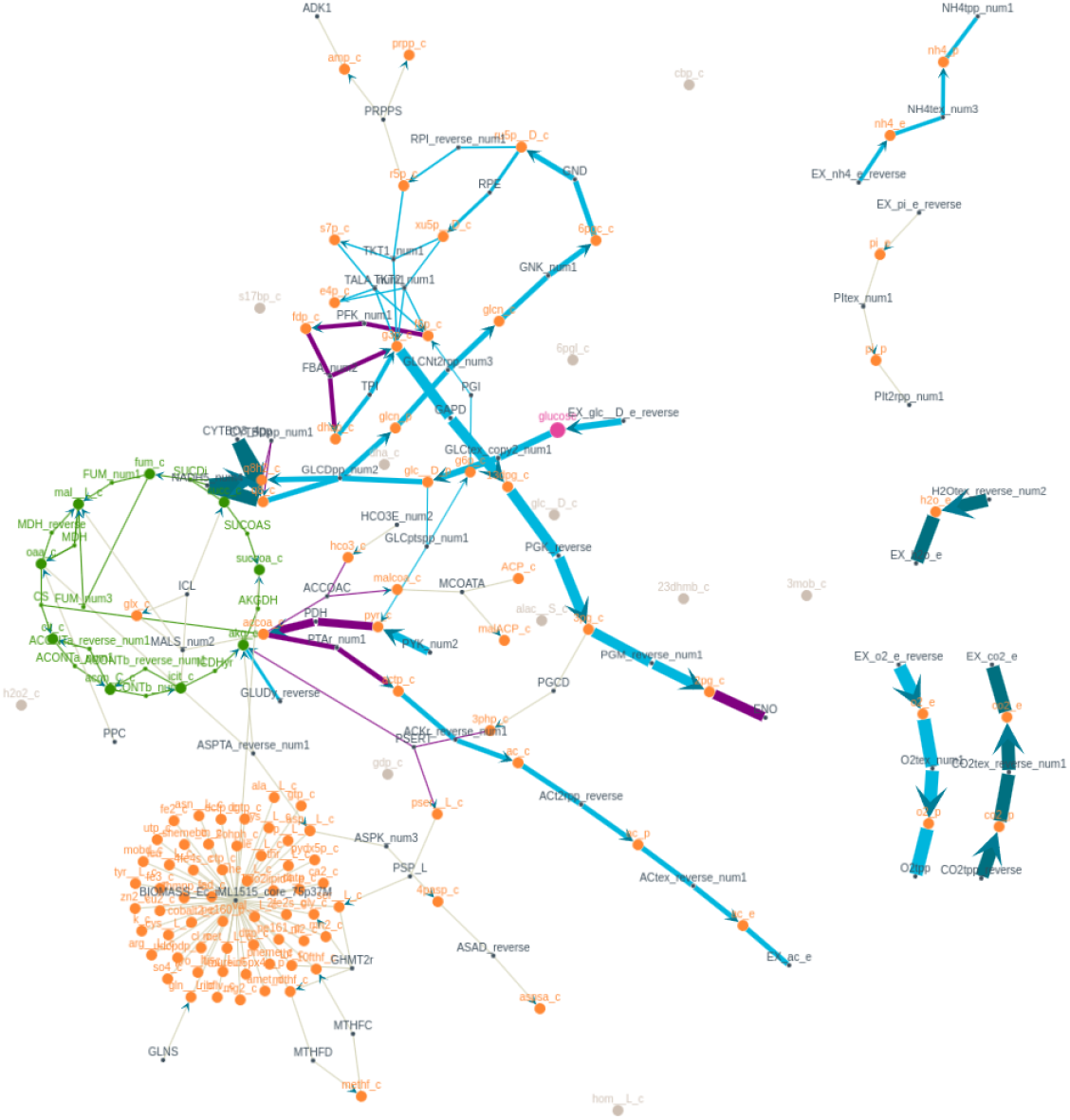
Metabolic Flux Visualisation: Enzyme-constrained solution, found with FBA for the min 8 clbr model.

## E. Proof of Equivalence of Flux Control Coefficient and Enzyme Cost

### E.1. Introduction

In an enzyme-constrained metabolic model, we have:

- A total enzyme budget *E*_*tot*_ that is distributed among different enzymes in the cell.
- Each reaction *v*_*j*_ limited by the amount of its enzyme *E*_*j*_ times the enzyme turnover number/catalytic rate *k*_*cat*_, *j*. That is,

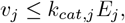

often under simplifing assumptions such as saturating conditions.
- An objective function (commonly biomass production) that we seek to maximise subject to stoichiometric and enzyme capacity constraints.

We want to *prove* that under optimal allocation of enzymes (i.e., at the solution maximising biomass flux *v*_*J*_ ),

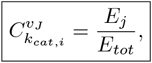

Where 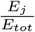 is the *fraction of total enzyme* allocated to enzyme *j* (the enzyme cost of that enzyme).

### E.2. Proof

#### E.2.1. Defining the Flux Control Coefficient

The flux control coefficient of reaction *j* with respect to *k*_*cat,j*_ is :

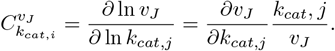

Here *v*_*J*_ is the biomass flux.

#### E.2.2. Total Enzyme Budget Constraint

Suppose we have a constraint on total enzyme:

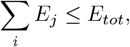

and we want to maximise *v*_*J*_. Because *v*_*j*_ = *k*_*cat,j*_*E*_*j*_, we can express

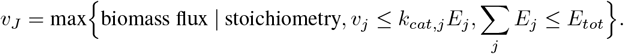

At the *optimal* solution, each active reaction satisfies *v*_*j*_ = *k*_*cat,j*_*E*_*j*_.

#### E.2.3. Lagrange Multipliers

We introduce a Lagrange multiplier *λ* for the total enzyme constraint. At the optimum:

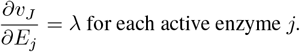

This condition states that each actively used enzyme must have the same marginal benefit; otherwise, shifting enzyme to higher-return enzymes could increase *v*_*J*_ .

#### E.2.4. Relationship Between *∂v*_*J*_ */∂k*_*cat,j*_ and *∂v*_*J*_ */∂E*_*j*_

Observe

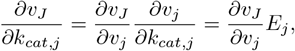

Because 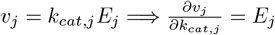.

Also,

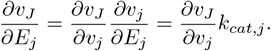

Given 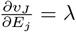, we get

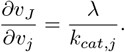

Hence,

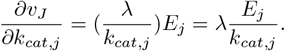

#### E.2.5. Express 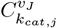 in Terms of *λ*

Recall

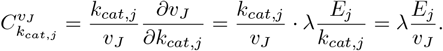

So

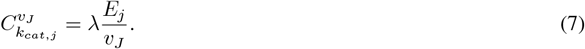

#### E.2.6. Summation Theorem and ∑_*j*_ *E*_*j*_ = *E*_*tot*_

Under typical control analysis assumptions (each *k*_*cat,j*_ is an independent parameter), the sum of the control coefficients 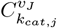 for all active *j* is 1:

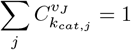

Combined with equation 7:

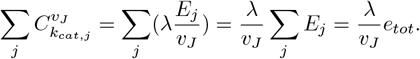

Because this sum equals 1, we obtain

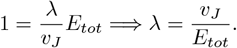

#### E.2.7. Final Step: Fraction of Total Enzyme = Flux Control Coefficient

Substituting 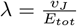 back into equation 7,

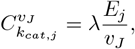

we get

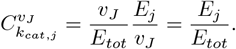

Hence,

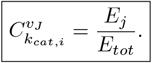

This completes the proof that, under an optimal solution (maximising flux with a fixed enzyme budget), the fraction of total enzyme allocated to each enzyme (the enzyme cost) is numerically equal to its flux control coefficient with respect to *k*_*cat,j*_. Equivalently, enzyme cost *∝* control coefficient.

## F. Enzyme abundances benchmarking

One of the benchmarks in our benchmarking process traces accuracy in enzyme abundances. Corrao et al. used polypeptide abundances from Schmidt et al. (2016) to compute enzyme abundances for 290 enzymes in *E. coli* (Corrao et al., 2024). We use their data for growth on glucose.

As the enzymes and reactions in their model have different names to ours, we matched our reactions to theirs by 1) finding all the genes for each enzyme in their model by using their knowledge graph and 2) finding all reactions in our model that use this specific set of genes. As some enzymes may catalyse multiple reactions, we first find the ‘abundance’ of each enzyme-reaction pair:

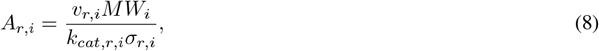

where the subscripts *r* and *i* denote reaction *r* catalysed by enzyme *i. MW*_*i*_ is the molecular weight of the enzyme. *A*_*r,i*_, *v*_*r,i*_, *k*_*cat,r,i*_, and *σ*_*r,i*_ are the abundance, flux, turnover number, and average saturation coefficient respectively. As mentioned in Section 3.1, throughout this work we assume a constant value of *σ*_*r,i*_ = 1. If the flux *v*_*r,i*_ was lower than the expected floating point accuracy of the solver (generously set to 10^−14^), the abundance for that reaction was set to zero, as it would be indiscriminately close to zero. To compute the total enzyme abundance for enzyme i, we then simply add the abundances for all reactions that this enzyme catalyses:

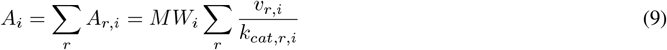

We then compare these abundances to the measured values from (Corrao et al., 2024). To do so, we follow (Corrao et al., 2024) in scaling the predicted abundances by a scale factor such that their sum is equal to the sum of all measured abundances ∑_*i*_ *A*_*i*_ = ∑_*i*_ *A*_*i,measured*_. We then compute the root mean squared logarithmic error (RMSLE; we use a logarithmic scale as the abundance values cover a range of 10^−12^ to 10^−1^).

## G Reference models

Here we present more detail about the reference models used in the benchmarking process in Section 4.2

- **ECMpy ML** is the ECMpy v1 model before doing any *k*_*cat*_ correction rounds, i.e. iML1515 irr enz constraint.json, produced by Mao et al. (2022) as per their GitHub on 25 Dec 2021.^6^ This model uses machine learning *k*_*cat*_ values from Heckmann et al. (2018).
- **ECMpy expmnt** is the ECMpy v2 model^7^ (Mao et al., 2024) before applying any *k*_*cat*_ corrections. Unlike the aforementioned ECMpy ML, which uses machine learning *k*_*cat*_s, the values in this model are experimental values. They are obtained from the BRENDA (Chang et al., 2021) and SABIO-RK (Wittig et al., 2018) databases using AutoPACMEN (Bekiaris & Klamt, 2020), and an average value of *k*_*cat*_ was chosen for reactions that were not in the databases.
- **ECMpy DLKcat** is a model created via the DLKcat pipeline in ECMpy v2 ^8^, which computes machine learning *k*_*cat*_s using the DLKcat method (Li et al., 2022). Unfortunately, the code in the ECMpy v2 GitHub produced NaN values for 15% of *k*_*cat*_s. This is almost entirely due to the fact that the code in the notebook does not retrieve the correct SMILES information for these reactions. We changed all NaN values to empty strings, so that these reactions were unconstrained. It must therefore be noted that since DLKcat could not be applied effectively to all reactions, our benchmarking results reflect the performance of the combined ECMpy+DLKcat pipeline rather than DLKcat alone.
- **ECMpy ML calibrated** is a model from ECMpy v1. This model, titled iML1515_irr_enz_constraint_adj_round1.json, is produced by subjecting the uncalibrated model ECMpy_ML to the first round of *k*_*cat*_ calibration. There is also a second correction round, which uses ^13^C fluxes, but we do not include this model as it uses additional experimental information and is therefore not comparable to our calibrated models. The first calibration round adjusted *k*_*cat*_s selected by enzyme cost 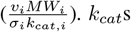 were updated for reactions that used more than 1% of total enzyme. The updated *k*_*cat*_s are pulled from the BRENDA and SABIO-RK databases. See Mao et al. (2022) for more details.
- **ECMpy expmnt calibrated** is the calibrated version of the ECMpy v2 model ECMpy expmt, which has been passed through the v2 *k*_*cat*_ calibration process. This process involved 50 rounds of iteratively adjusting *k*_*cat*_ for those enzymereaction pairs with the highest enzyme cost, and updating its value to the highest datapoint found in BRENDA and SABIO-RK. See Mao et al. (2024) for more details.

## H. *k*_*cat*_ sensitivity through flux control coefficients

**Figure 8.**
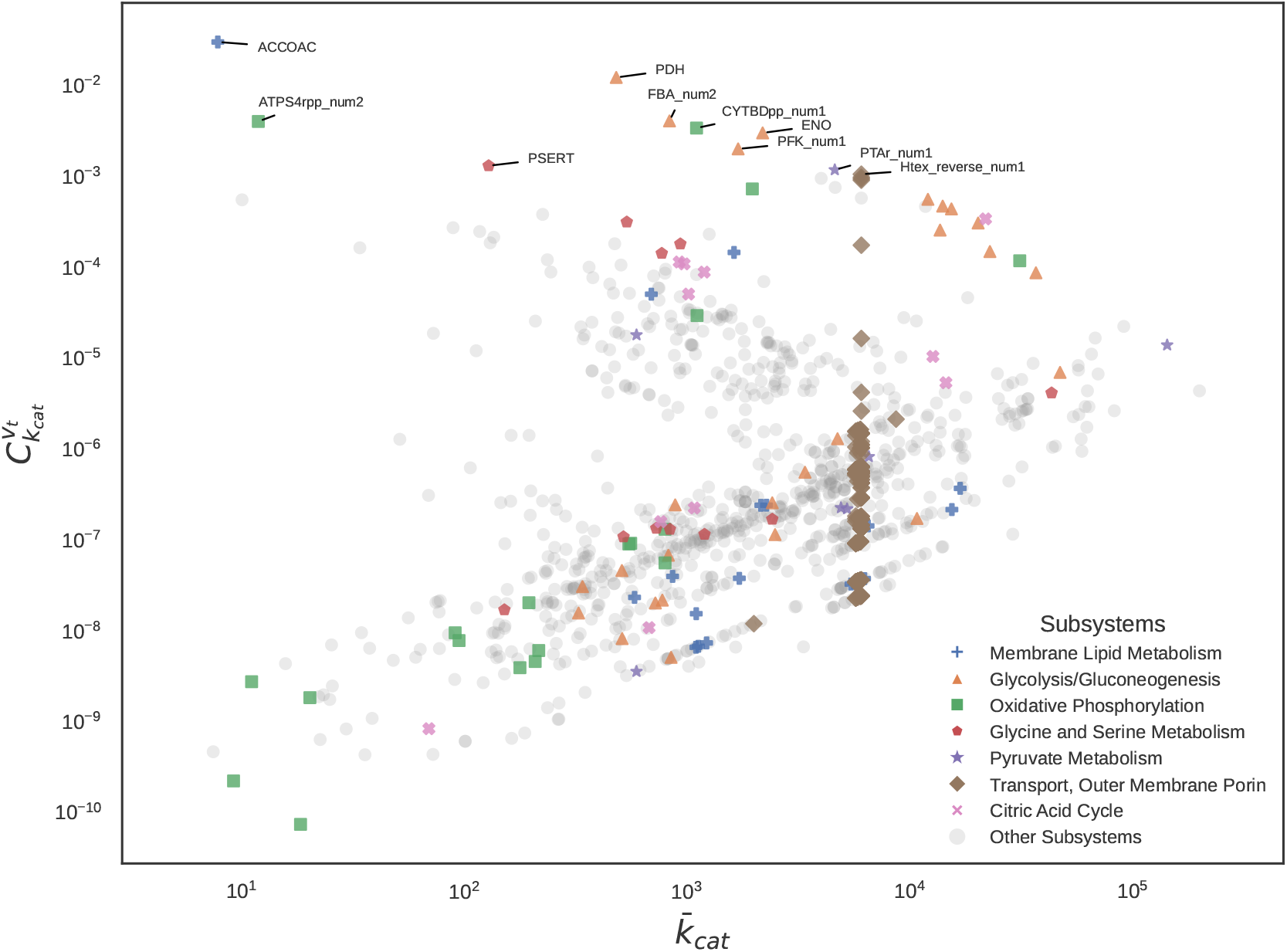
*k*_*cat*_ sensitivity plot, categorised and coloured by subsystem for the min model (subsystem annotations from the BiGG database (King et al., 2015)). Flux control coefficient 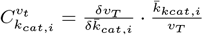, rescaled 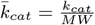, perturbation 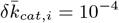, metabolic target *T* : *BIOMASS Ec iML*1515 *core* 75*p*37*M*. Coloured subsystems are those present in the top 10 most sensitive reactions under flux control coefficient analysis except the Citric Acid Cycle which is highlighted for reference. The band of Transport reactions at 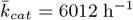 is due to the heuristic described in Section 3.3. Not shown are reactions where 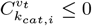 (due to log-scale plotting).

## I. Top 10 most sensitive reactions for min model by flux control coefficient

**Table 4.**
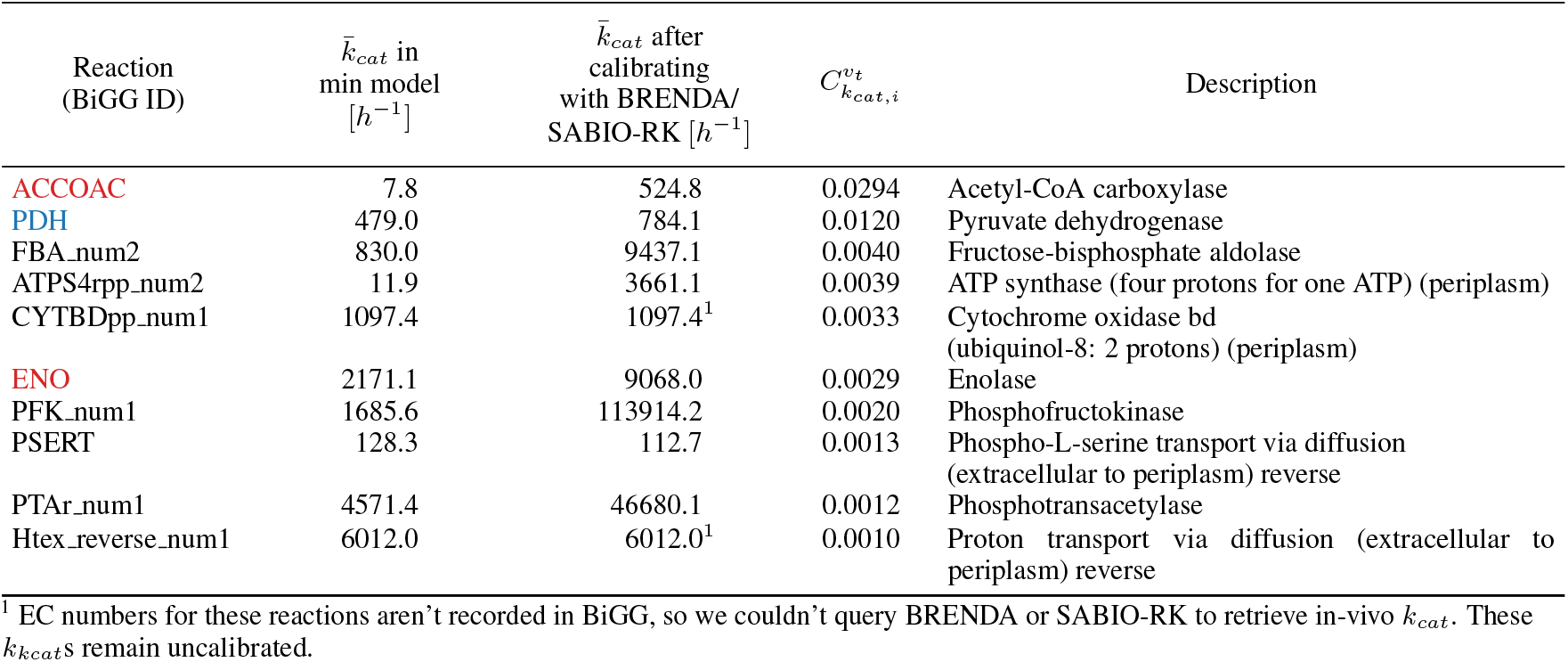
Top 10 most sensitive reactions for the min model as measured by flux control coefficient, 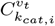. Red highlighting corresponds to ECMpy v1 round 1 enzyme abundance corrections, Blue highlighting corresponds to ECMpy v1 round 2 ^13^C corrections. EC Numbers for each reaction were queried across BRENDA and SABIO-RK, and we retrieved the largest *k*_*cat*_ across each search (including reactions from different species to *E. coli*, and across different environmental conditions).

| Reaction<br>(BiGG ID) | $\bar{k}_{cat}$ in<br>min model<br>[$h^{-1}$] | $\bar{k}_{cat}$ after<br>calibrating<br>with BRENDA/<br>SABIO-RK [$h^{-1}$] | $C_{k_{cat},i}^{v_t}$ | Description |
| --- | --- | --- | --- | --- |
| ACCOAC | 7.8 | 524.8 | 0.0294 | Acetyl-CoA carboxylase |
| PDH | 479.0 | 784.1 | 0.0120 | Pyruvate dehydrogenase |
| FBA_num2 | 830.0 | 9437.1 | 0.0040 | Fructose-bisphosphate aldolase |
| ATPS4rpp_num2 | 11.9 | 3661.1 | 0.0039 | ATP synthase (four protons for one ATP) (periplasm) |
| CYTBDpp_num1 | 1097.4 | 1097.4<sup>1</sup> | 0.0033 | Cytochrome oxidase bd<br>(ubiquinol-8: 2 protons) (periplasm) |
| ENO | 2171.1 | 9068.0 | 0.0029 | Enolase |
| PFK_num1 | 1685.6 | 113914.2 | 0.0020 | Phosphofructokinase |
| PSERT | 128.3 | 112.7 | 0.0013 | Phospho-L-serine transport via diffusion<br>(extracellular to periplasm) reverse |
| PTAr_num1 | 4571.4 | 46680.1 | 0.0012 | Phosphotransacetylase |
| Htex_reverse_num1 | 6012.0 | 6012.0<sup>1</sup> | 0.0010 | Proton transport via diffusion (extracellular to<br>periplasm) reverse |

<sup>1</sup> EC numbers for these reactions aren't recorded in BiGG, so we couldn't query BRENDA or SABIO-RK to retrieve in-vivo $k_{cat}$ . These $k_{k_{cat}}$ s remain uncalibrated.

## J. Top 100 most sensitive reactions according to flux control coefficient

**Figure 9.**
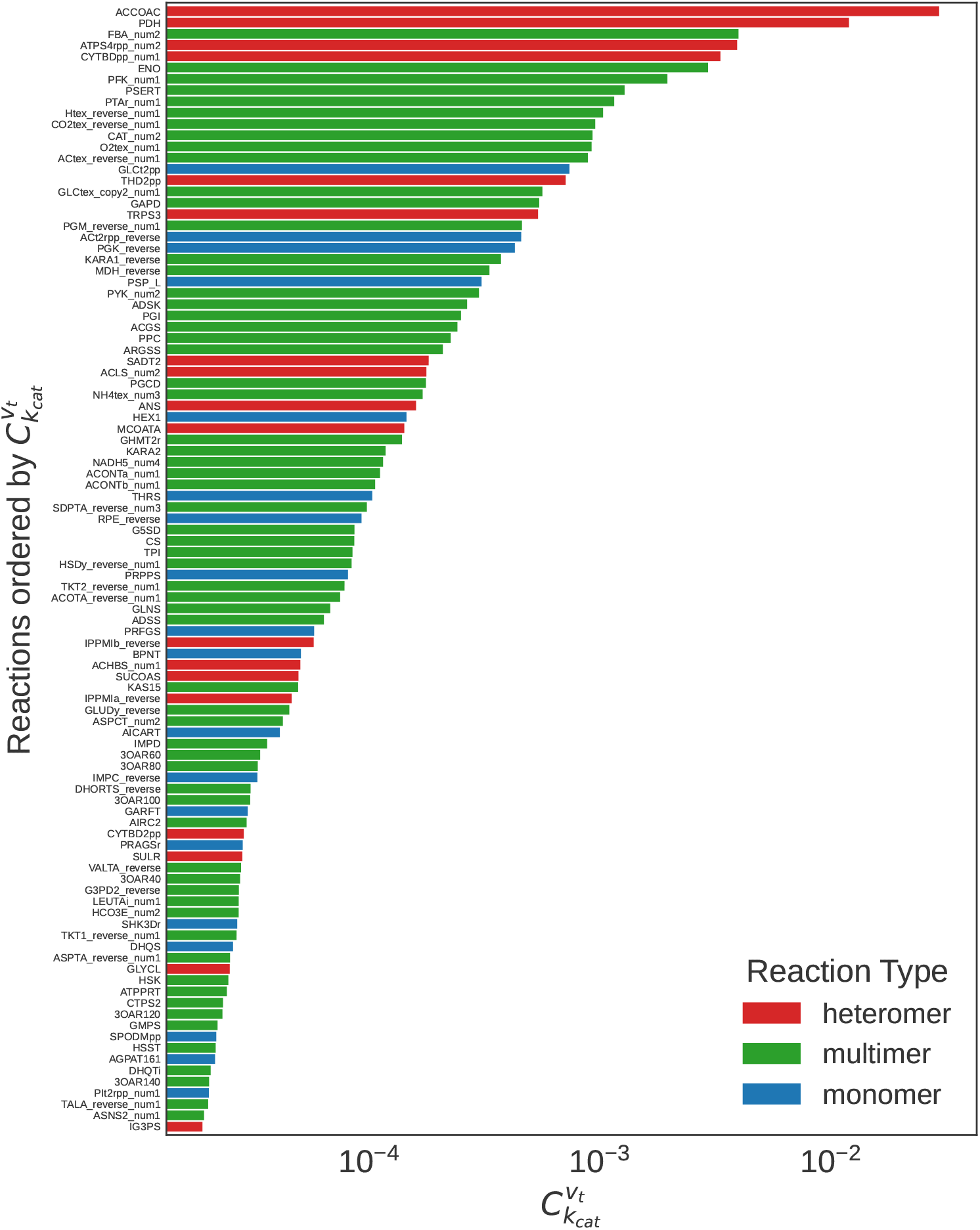
Top 100 reactions in the min model ranked by flux control coefficient 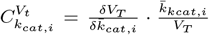, coloured by enzymatic reaction type (heteromer, homomultimer, monomer). Non-enzymatic reactions are not shown.

## Footnotes

1 Often there are a large number of reactions for which no *k*_*cat*_s are available and one needs to generate these in an ad-hoc fashion.

2 This treatment of *p*_*tot*_ and *f* assumes constant protein mass for enzyme production across different conditions (e.g., media), a simplification which may not hold true in real-life scenarios.

3 There is a question around whether the true turnover rate would be higher than this, since evolutionarily speaking, the mean reason for forming a multimer would be to reduce the reaction’s activation energy, but addressing this issue is beyond the scope of our work.

4 We only perturb a single *k*_*cat*_ at a time, as combinations of perturbations would lead to combinatorial explosions.

5 Some reactions are missing an associated gene in the underlying iML1515 model. Some of these reactions are COBRA boundary reactions of the exchange (EX ) type and are essentially spontaneous reactions, but not all of them. To be on the safe side, all such reactions were not annotated, and did not receive a *k*_*cat*_ value in our model (see Appendix B.

6 http://github.com/tibbdc/ECMpy/tree/433463a9b22994765351eae1ea1b74d133f7a483

7 To be precise, we create this model by running the notebook “02.get ecModel using ECMpy.ipynb” on the saved state of the ECMpy GitHub repository as it was on 15 Feb 2025 (https://github.com/tibbdc/ECMpy).

8 As this model is not saved to their Github we had to generate it ourselves. We did this via the notebook 01.get reactiion kcat using DLKcat [sic]. We also had to change line 4 in cell 6 to `subbnumdf = pd.read_csv(gene_subnum_path, index_col = 0)`; without this change the pipeline does not compute *k*_*cat*_*/MW* correctly.

